# Reshuffling of the coral microbiome during dormancy

**DOI:** 10.1101/2022.08.16.504223

**Authors:** Anya L Brown, Koty Sharp, Amy Apprill

## Abstract

Quiescence, or dormancy, is a response to stressful conditions in which an organism slows or halts physiological functioning. Although most species that undergo dormancy maintain complex microbiomes, there is little known about how dormancy influences and is influenced by the host’s microbiome, including in the temperate coral, *Astrangia poculata*. Northern populations of *A. poculata* undergo winter quiescence. Here, we characterized wild *A. poculata* microbiomes in a high-resolution sampling time series before, during, and after quiescence using 16S ribosomal RNA gene sequencing on active (RNA) and present (DNA) microbiomes. We observed a restructuring of the coral microbiome during quiescence that persisted after re-emergence. Upon entering quiescence, corals shed copiotrophic microbes, including putative pathogens, suggesting removal of these taxa as corals cease normal functioning. During and after quiescence, bacteria and archaea associated with nitrification were enriched, suggesting the quiescent microbiome may replace essential functions through supplying nitrate to corals and/or microbes. Overall, this study demonstrates that key microbial groups related to quiescence in *A. poculata* may play a role in the onset or emergence from dormancy, and long-term regulation of the microbiome composition. The predictability of dormancy in *A. poculata* provides an ideal natural manipulation system to further identify factors that regulate host-microbial associations.

**Importance:** Using a high-resolution sampling time series, we are the first to demonstrate a persistent microbial community shift with quiescence (dormancy) in a marine organism, the temperate coral, *Astrangia poculata.* Furthermore, during this period of community turnover, there is a shedding of putative pathogens and copiotrophs and an enhancement of the ammonia-oxidizing bacteria (Nitrosococcales) and archaea (*Ca*. Nitrosopumilus). Our results suggest that quiescence represents an important period during which the coral microbiome can “reset,” shedding opportunistic microbes and enriching for the re-establishment of beneficial associates, including those that may contribute nitrate while the coral animal is not actively feeding. We suggest this work provides foundational understanding of the interplay of microbes and the host’s dormancy response in marine organisms.

## Introduction

Nearly all animal, plant and bacterial phyla include species that undergo dormancy to survive periods of harsh environmental stress. Dormancy represents a resting state, in which metabolic functions are depressed (Hand, 1991; Cáceres, 1997). In bacteria, dormancy ensures their persistence in hosts, and is a trait of both pathogens and beneficial symbionts (Flint *et al*., 2005; Barry *et al*., 2009). In general, dormancy can be composed of multiple phases: preparation (before dormancy begins); initiation (onset of dormancy); maintenance (metabolic suppression, depletion of energy stores); potentiation (beginning of the post-dormant periods); and activation (resume activity, Wilsterman *et al*., 2021).

Hosts that undergo dormancy are also involved in complex associations with microorganisms. Host-associated microbes are involved in host immunity, physiology, survival, and metabolic function, thus likely are influenced and can be influenced by dormancy. The role of microbes in host dormancy is an emerging field, thus our understanding of the microbial role, and even our understanding of the microbial shifts surrounding dormant periods are limited. Indeed, microbes may influence the onset and cessation of dormancy, or replace host functioning during periods of dormancy.

In several hosts, including bears, squirrels, crickets and parasitoid wasps (Carey *et al*., 2013; Sommer *et al*., 2016; Dittmer and Brucker, 2021; Regan *et al*., 2022), dormancy is associated with shifts in the composition of the host’s microbiome. One role these community shifts play may be to replace resource acquisition or use while host functioning is shut down or reduced (Carey *et al*., 2013; Sommer *et al*., 2016; Dittmer and Brucker, 2021; Regan *et al*., 2022). For example, in ground squirrels, the restructuring of the gut is mediated by food availability (Carey *et al*., 2013). During hibernation the gut microbiome plays an important role in nitrogen recycling, while the squirrel is fasting (Regan *et al*., 2022). Dormant states are also associated with pathogen avoidance, for example, nematodes enter diapause to avoid infection (e.g., by not ingesting pathogens; (Palominos *et al*., 2017).

In aquatic invertebrates, the onset of dormancy, or quiescence, is associated with harsh environmental conditions, such as winter (Cáceres, 1997). Few examples of dormancy are found in cnidarians, and even fewer in the class Anthozoa. However, the temperate scleractinian coral, *Astrangia poculata,* is known to undergo quiescence in the winter months, which is thought to be a response to extreme cold temperatures (Grace, 2017). Similar to other species that undergo dormancy, quiescent *Astrangia poculata* have a distinct phenotype. They pull in their tentacles, form a puffed-up ring around their oral disc, do not respond to tactile stimulation, and do not actively feed. During quiescence there are also physiological shifts including lowered coral growth rates (Grace, 2017), polyp loss (Dimond *et al*., 2013), and shifts in the coral transcriptome, associated with thermal stress and lowered motility (Wuitchik *et al*., 2021). Additionally, the physiological costs of dormancy can last beyond winter into spring (Trumbauer *et al.*, 2021).

*Astrangia poculata* represents a multi-domain symbioses, involving specific bacteria and archaea (Sharp et al., 2017), and it engages in facultative symbiosis with the eukaryotic microalga *Breviolum psygmophilum* (Family Symbiodiniaceae), the same genus of microalgae as those found in many tropical corals (Thornhill *et al*., 2008). This species shows two forms: a “white” or aposymbiotic phenotype and a “brown” or symbiotic phenotype, depending on visible presence of microalgae in their otherwise transparent tissues. Although in symbiosis with photosynthetic algae, the coral mainly relies on heterotrophy for nutrition (Dimond and Carrington, 2007; Trumbauer *et al*., 2021).

*A. poculata* microbiomes are dominated by taxa similar to tropical corals at the class level (e.g., γ- and α proteobacteria; Cytophagia, Flavobacteria), although the *A. poculata* microbiome is generally less diverse (Sharp *et al*., 2017). As the similarities in taxa suggest, the microbiome of *Astrangia* also is expected to function similarly to tropical corals: in nutrient cycling, sources of nutrition, immunity, and defense (Bourne *et al*., 2016; Peixoto *et al*., 2017; Robbins *et al*., 2019; Rosado *et al*., 2019; Santoro *et al*., 2021).

*A. poculata* microbiomes shift with season. The *A. poculata* winter microbiome is enriched in Clostridiaceae, Flavobacteriaceae, and Rickettsiaceae (Sharp *et al*., 2017). In the spring, the microbiome alters in composition to a less diverse and less variable microbial community compared to winter, fall and summer (Sharp *et al*., 2017). The shift in microbial communities from winter to spring also corresponds to tropical coral microbiomes that undergo cyclical mucus shedding (Glasl *et al*., 2016). The seasonal shifts in *Astrangia* microbiomes are thought to be associated with quiescence, however, a detailed characterization of the microbial shifts that occur around quiescence is needed to determine how dormancy may impact the microbiome, and vice-versa.

Here, we collected a high-resolution sampling time series to characterize the shift in microbiome diversity and community structure as *Astrangia poculata* corals go into, remain in, and come out of quiescence (Fig 1). Based on the results of seasonal studies and other studies on animal dormancy, we expected a shift in community composition throughout quiescence, lowered diversity of microbes in the winter, and decreased variability among individual coral colonies as they emerged from quiescence. As some microbes may also be dormant as the coral host enters dormancy, we compared the active (RNA) and present (DNA) microbiome over time, to understand which taxa are contributing to host-associated microbiome activity during host dormancy. Lastly, we propose new hypotheses about the taxonomic shifts, their functional significance, and the implications on the host throughout the phases of dormancy (before, during, after).

**Figure 1.**
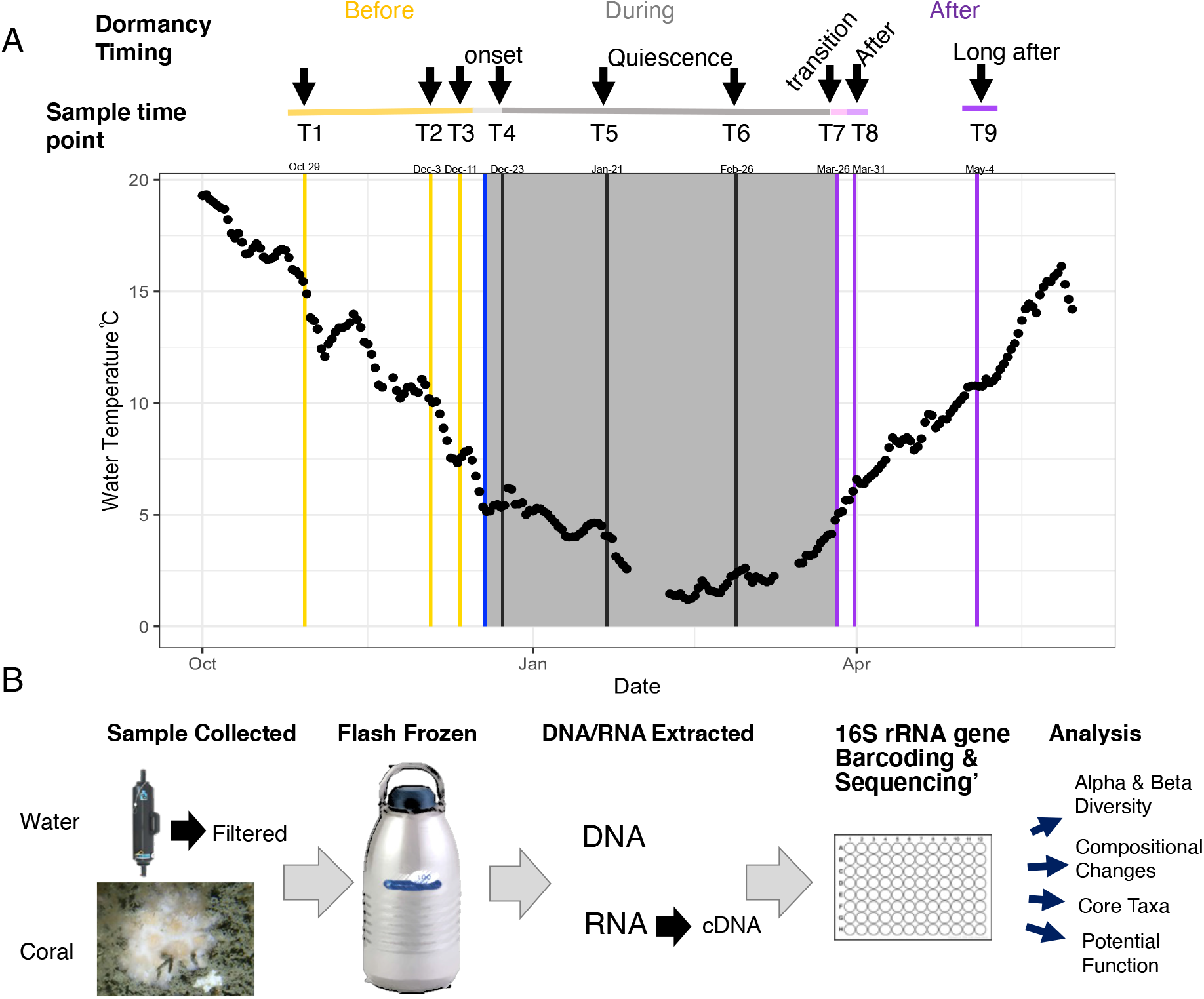
Overview of field experimental sampling, sample processing and analysis. (A) Mean daily temperature based on the Station BZBM3 in Woods, Hole MA. Dates range from October 2020-May 2021. The shaded region represents the time in which corals were in quiescence. Lines refer to sampling periods around dormancy: yellow indicates “before”, the blue line refers to when corals went into quiescence; gray lines refer to quiescence; purple lines refer to “after” quiescence. Above the plot, the line and labels refer to when samples were taken, the designation of sample points, and naming of the phases of dormancy. (B) Schematic of the sampling protocol for corals (n = 10 per timepoint) and water (n = 4 per timepoint) and analyses.

## Methods

### Sample Collection

From late October 2020-May 2021 we collected 10 distinct “white” or aposymbiotic colonies of *A. poculata* at each of nine time points (Table 1, Figure 1). Corals were collected on SCUBA at 18m from pilings on the Woods Hole Oceanographic Institution’s Iselin dock (41°31’25.1”N 70°40’19.3”W). We selected aposymbiotic colonies to reduce the potential effects of the algal symbionts in our understanding of dormancy-related microbial shifts. Selected colonies showed no visible coloration in any of the polyps (or near absence of algae; Figure 1). Corals were collected using a hammer and chisel and were frozen in liquid nitrogen vapors immediately upon surfacing from the dive.

**Table 1.** Coral and seawater sampling dates, time points, and designation of quiescence phases for different analyses.

| Date | Temperature C | Time Point | Timing (NMDS) | Dormancy timing |
| --- | --- | --- | --- | --- |
| 29-October-2020 | 15.1 | T1 | Before | Before |
| 3-December-2020 | 10.2 | T2 | Before | Before |
| 11-December-2020 | 7.3 | T3 | Before | Before |
| 23-December-2020 | 5.3 | T4 | Onset | During |
| 21-January-2021 | 4.1 | T5 | During | During |
| 26-February-2021 | 2.3 | T6 | During | During |
| 26-March-2021 | 4.8 | T7 | Transition | During/After (depending<br>on the coral |
| 31-March-2021 | 6.1 | T8 | After | After |
| 4-May-2021 | 10.8 | T9 | Long after | After |

During eight of the time periods (starting on 3-Dec), we collected water samples via a 5 L Niskin bottle triggered at depth (18 m). We subsampled this water for inorganic nutrients (25 ml, frozen to −20 °C), which were analyzed as described previously (Weber *et al*., 2020). For seawater microbial community analysis, we filtered 500 ml of the collected water through a 0.22 μm Sterivex filter (Millipore Sigma, Burlington, MA; n=4 per time point) using peristaltic pressure, and then flash froze the filter in liquid nitrogen vapors.

### Temperature data

Seawater temperature data was used from Station BZBM3 in Woods, Hole MA, measured 1.7m from mean lower low water line (NDBC Station History Page, 1996). Temperature data was averaged by day over the time period of sampling (October-May).

### Nucleic Acid Extractions and cDNA Synthesis

For both coral and water samples, DNA and RNA were extracted using the Quick DNA/RNA mini prep plus kit with ZR BashingBeads (0.1 and 0.5mm; Zymo Research, Carlsbad, CA). For coral samples, one polyp (including mucus, tissue and skeleton) of each coral colony was removed with a sterilized chisel and hammer. Coral fragments were added to the bead tubes and then suspended in 800 μl of DNA/RNA Shield and bead beat for 10 min at top speed on a vortexer. We added proteinase K (15 μl of 20 mg μl^-1^) to further breakdown cells and isolate the DNA, and then continued with the rest of the steps in the manufacturer’s protocol including the DNAase step for the RNA portion of the sample. Extracted RNA was frozen at −80 °C, and DNA was frozen in a −20 °C until further analysis.

For water samples, we opened the plastic case of the sterivex filter using a sterilized steel cutting implement and added 1000 μl of DNA/RNA shield to the sample and bead tube. Then, 30 μl of proteinase K (20 mg μl^-1^) was added before following the manufacturers protocol. Extraction blanks, which included reagents but no samples (n =3 for water protocol, n = 6 for the coral protocol), were carried out as well for both RNA and DNA.

We converted RNA to cDNA for sequencing the active microbiome using the New England Biolabs (Ipswich, MA) ProtoScript^®^ II First Strand cDNA Synthesis Kit. We followed the standard protocol and used 2 μl of coral RNA and 6 μl of water RNA as template.

### 16S rRNA gene Library Prep

We prepared DNA and cDNA for 16S rRNA gene sequencing of the V4 region using barcoded 515FY (Parada *et al*., 2016) and 806RB (Apprill *et al*., 2015) primers that target bacteria and archaea with standard barcodes (Kozich *et al*., 2013). The PCRs (25 μl) were performed in duplicate per sample, and prepared using the High Fidelity Phusion Master Mix with HF buffer (12.5 μl /sample), DMSO (0.75 μl /sample) (New England Biolabs), molecular grade water (7.25 μl /sample), the primers (1.25 μl of each), and 1 μl of template. The thermocycler conditions were: an initial denaturation step of 95 °C for 2mins, and then 30 cycles of 95°C (20s), 55°C (15s), 72°C (5min), and a final elongation step of 72 °C for 10 min.

Each PCR reaction was run out on a 1.5% agarose, and the correct band (determined by the location of the positive control) was excised. Excised bands were extracted and purified with the MinElute Gel Purification kit (Qiagen Inc., Germantown, MD). Purified PCR products were quantified with a Qubit, diluted to 1 ng μl^-1^, and then pooled at 5 ng of purified product per sample. Each pool contained negative PCR controls with no visible bands, and a Mock community (Even, low community B; BEI Resources). The pools (3 total) were sequenced on an Illumina MiSeq, with 250bp paired-end sequencing.

### Bioinformatics

With the de-multiplexed forward and reverse sequences, we used the DADA2 pipeline (Callahan *et al*., 2016) in R (R Core Team, 2021) for quality control, merging sequences and assigning Amplicon Sequence Variants (ASVs). Forward and reverse reads were visually inspected for quality and to determine the cut off values in the filter and trim step: filterAndTrim(fnFs, filtFs, fnRs, filtRs, truncLen = c(240, 150), maxN = 0, maxEE = c(2,2), rm.phix = TRUE, compress = TRUE, multithread = TRUE). Error rates were computed, and used for sequence inference. Sequences were then merged and ASV tables created. Because of the size of the dataset, error rates and ASV tables were created per MiSeq run, the tables were then merged, and chimeras were checked and removed. Taxonomy was assigned using the Silva v132 training set (Quast *et al*., 2013; Yilmaz *et al*., 2014), and retrieval of taxa from mock communities was checked.

The taxa table, ASV table and metadata table were loaded into phyloseq (McMurdie and Holmes, 2013), where chloroplasts and mitochondria were removed. Using the decontam package (Karstens *et al*., 2019), we removed contaminant taxa using the prevalence of taxa (at 0.01) in the negative controls (including PCR negatives, extraction kit blanks, water filter blanks for both DNA and cDNA).

Alpha diversity was calculated Hill numbers D^0^, D^1^, and D^2^, which correspond to richness (rarefied), exponentiated Shannon diversity and the Inverse Simpson index respectively (Alberdi and Gilbert, 2019). Higher D values indicate more even and speciose communities. We estimated diversity indices using phyloseq.

To compare variability in coral communities over time, we computed Bray Curtis dissimilarities on the relative abundances of taxa within a sample. We then quantified the distance from each sample point to the group’s centroid (betadispersion) using the betadisper function in the vegan package (Oksanen *et al*., 2016).

We tested for significant differences in alpha diversity and dispersion using linear models, comparing dormancy timing (before, during, after), sample time (1-9) and active/present microbiome (cDNA/DNA) using linear models in R (R Core Team, 2021). Residuals were visually inspected to meet assumptions of heteroscedasticity and normality. Significance was assessed using Anova from the car package ((Fox J, 2019). When necessary, post hoc tests (Tukey HSD) were used to evaluate differences among levels in treatments.

To examine compositional changes within *Astrangia poculata*’s microbiome throughout the phases of dormancy (before, at the onset, during, and after), we used Bray Curtis dissimilarity matrices based on relative abundance. We then compared the microbial community composition using PERMANOVA in the vegan package (Oksanen *et al*., 2016) with sample times (1-9) and the timing around dormancy (before, during, after) and active/present microbiome as factors. To understand the differences in community composition across time points and type of sample, we used pairwise comparisons with the EcolUtils package (Salazar 2022). Coral and water microbial communities were visualized on an NMDS plot.

We assessed which microbial taxa change in relative abundance with respect to timing of dormancy (before versus during, before versus after) using corncob (Martin *et al*., 2020). This method uses a beta-binomial model on the counts (number of reads) for each ASV to determine which taxa are significantly enriched from a reference level (in this case, before dormancy) in different treatments, and compares them iteratively. We compared the active (DNA) and present (RNA) microbiomes from the coral and water separately.

To determine which microbial taxa are associated with the “core” of *Astrangia poculata*, and how these taxa change over time, we determined which taxa were present at 80% prevalence across all samples within a time point (Bourne *et al*., 2016) with the microbiome package (Lahti and Sudarshan, 2017).

We inferred function of the microbiome using a functional inference tool based on taxonomy. Although there are drawbacks to tools that predict function from taxonomy (Lan *et al.*, 2016; Escobar-Zepeda *et al*., 2018), here, we took a conservative approach and used broad categorizations of functions using the FAPROTAX database (Louca *et al*., 2016) and microeco package (Liu *et al*., 2022) in R, which were generated from published metabolic and ecological functions and suggested for environmental data (Liu *et al*., 2021). We extracted the functions associated with ASVs that were determined to be significantly enriched based on the corncob analysis to understand potential functional shifts across the dormancy time periods in the active and present microbiomes.

## Results

*Astrangia poculata* collections occurred via SCUBA in Woods Hole, MA over a six-month period and began in late October when seawater temperatures were 15.4°C, during which time the corals polyps were extended and presumed to be feeding and metabolically active (Table 1, Fig 1A). Three collections (each of 10 colonies) occurred during this pre-quiescence period (Timepoints T1-3). Corals (at 16 m) were observed to be quiescent on Dec 18 (5°C) by divers who observed the area daily. At this time, polyps were retracted and presumed to be no longer feeding. On Dec 23, 2020, the first quiescent corals were collected (5°C) (T4; *n* = 10), and collections continued during quiescence (T5 and T6, *n* = 10 colonies each). On 24 Mar (5°C; T7), some corals emerged from quiescence, and some did not, five quiescent and five emerged corals were collected. By 31 Mar (T8), all corals had emerged from quiescence (n = 10), and collections continued for one additional post-quiescence period (15 April; T9; n = 10).

To investigate the microbial community associated with the coral and at each timepoint, the 16S rRNA genes of bacteria and archaea were amplified from DNA (present microbiome) and cDNA (active) extracted from one polyp of each coral colony and sequenced. Bacteria and archaea sequences were also obtained from seawater adjacent to the coral habitat (*n* = 4 per timepoint) using the same approach (Fig 1B). Raw sequences can be found in the NCBI SRA database (Bioproject: PRJNA860933).

After quality filtering and removal of taxa associated with the controls (0.73% of all ASVs), and chloroplasts and mitochondria (6.7% of unique ASVs) we retained 12,964,163 sequences (median = 30029.5) across all 238 samples (coral and water) and 19,656 unique ASVs. Four cDNA coral samples from different time periods (T1, T3, T4, T5,) were removed because of low number of sequences (<1000). Unique ASVs were examined per sample time, and in the coral present and active microbiome, we observed 10,504 and 9,681 unique ASVs, respectively; in the water present and active microbiome we observed 1,184 and 6,060 unique ASVs, respectively. After rarefying (only used in the Hill number D^0^ or richness analysis), there were 1,307 sequences/sample and 8242 unique ASVs across the dataset (water and coral).

### Alpha Diversity

As expected, there were fewer taxa in the active coral microbiome compared to the present coral microbiome, this was particularly evident in rarefied richness during and after dormancy (Fig 2A, Table 2: Active/Present Microbiome < 0.05 for all diversity measures). Interestingly, for active microbiomes, we observed a significant decrease in alpha diversity as corals went into dormancy, remained low as corals were in dormancy, and then began to increase as corals exited dormancy (Dormancy Timing in D^0^ or richness: p= 0.004; and D^1^or exponentiated Shannon diversity: p = 0.002; & Active/Present Microbiome <0.05, Fig 2 A,B: Table 2). However, in the present microbiome, there were no significant differences in diversity between before and during dormancy, but similarly, we observed an increase after dormancy based on TukeyHSD tests (in D^0^ and D^1^, Fig 2 A,B). Hill number D^2^ or the inverse Simpson Index, the diversity measure influenced by dominance, only significantly increased after corals came out of quiescence in both the active and present microbiomes (Dormancy timing: p = 0.002; Fig 2C).

**Figure 2.**
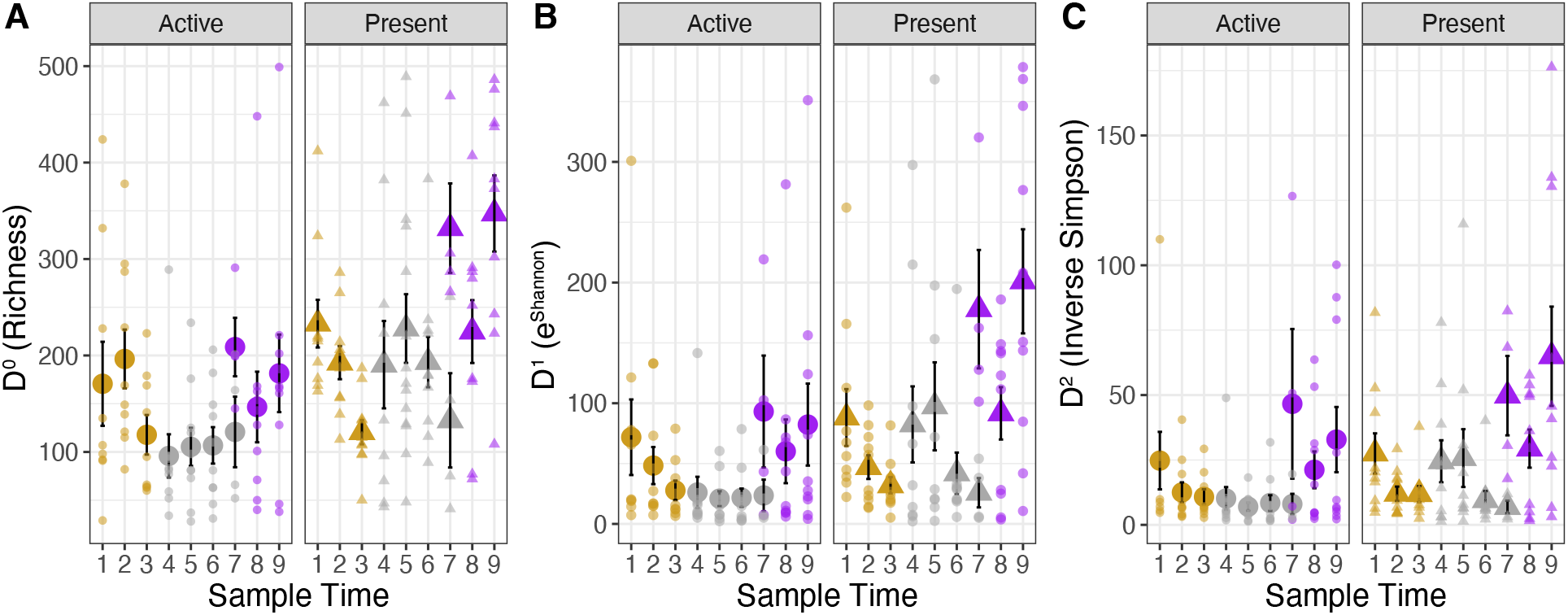
Coral microbiome alpha diversity decreases during quiescence. Plots show the mean ± se of Alpha Diversity measures. Colors indicate before (yellow), during (gray) and after (purple) quiescence. The active microbiome and present microbiome are separated by facets and by shape (circles and triangle, respectively). Smaller, transparent points represent raw values. A) Hill D^0^ (Rarefied richness), B) D^1^ exponentiated Shannon diversity (not rarefied), C) D^2^ or Inverse Simpson diversity. D^0^ and D^1^ decreased during quiescence, and then increased after quiescence in active microbes, however, in the present microbes, the diversity only changed (increased) after dormancy. Conversely, D^2^ remained low before and during quiescence and increased after quiescence for both DNA and cDNA.

**Table 2.**
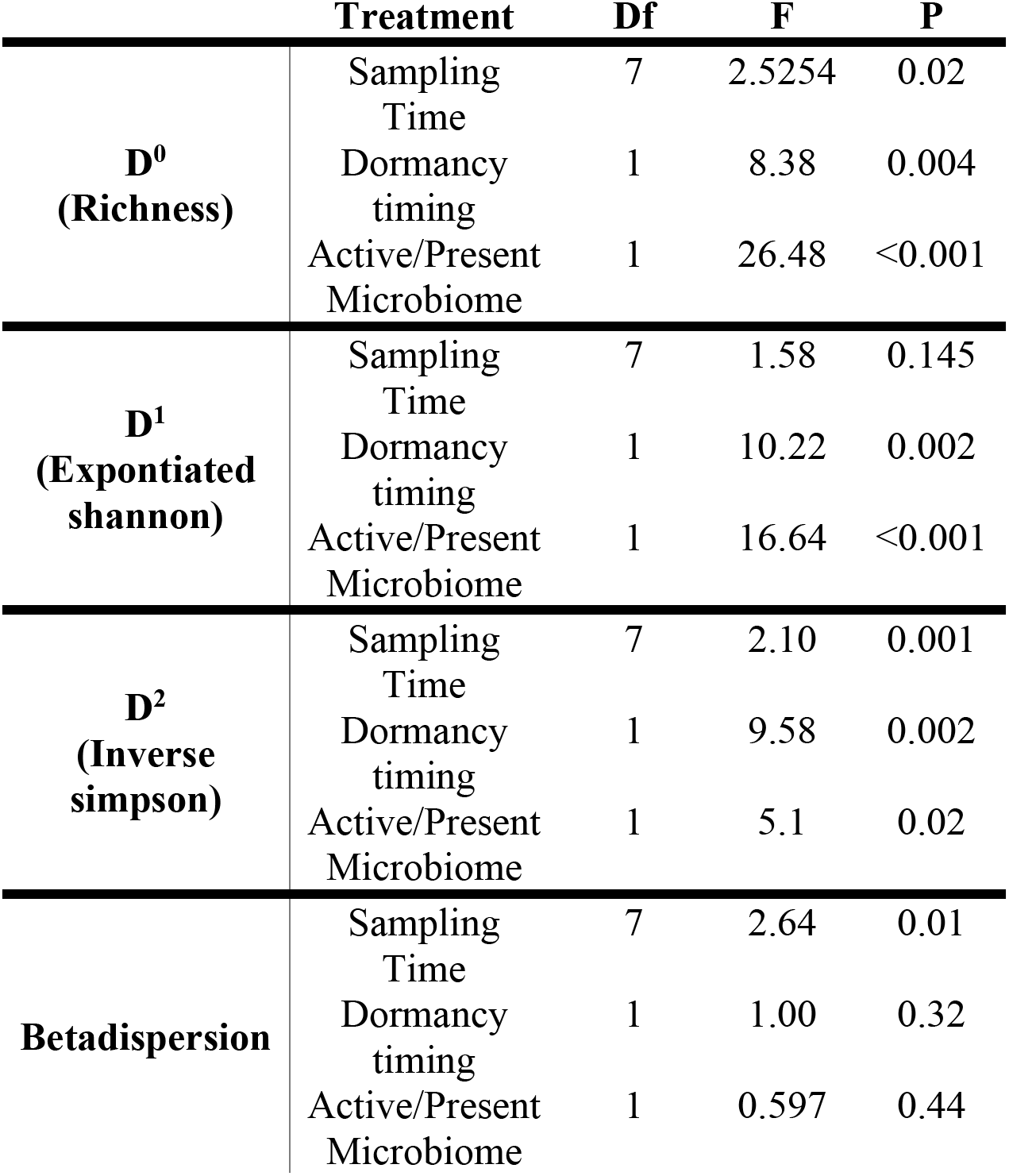
Results of statistical analysis examined for diversity indices, from ANOVA analysis. DF = degrees of freedom, F = frequency, P = p-value.

|  | **Treatment** | **Df** | **F** | **P** |
| --- | --- | --- | --- | --- |
| **D<sup>0</sup>**<br>**(Richness)** | Sampling Time | 7 | 2.5254 | 0.02 |
|  | Dormancy timing | 1 | 8.38 | 0.004 |
|  | Active/Present Microbiome | 1 | 26.48 | <0.001 |
| **D<sup>1</sup>**<br>**(Exponentiated shannon)** | Sampling Time | 7 | 1.58 | 0.145 |
|  | Dormancy timing | 1 | 10.22 | 0.002 |
|  | Active/Present Microbiome | 1 | 16.64 | <0.001 |
| **D<sup>2</sup>**<br>**(Inverse simpson)** | Sampling Time | 7 | 2.10 | 0.001 |
|  | Dormancy timing | 1 | 9.58 | 0.002 |
|  | Active/Present Microbiome | 1 | 5.1 | 0.02 |
| **Betadispersion** | Sampling Time | 7 | 2.64 | 0.01 |
|  | Dormancy timing | 1 | 1.00 | 0.32 |
|  | Active/Present Microbiome | 1 | 0.597 | 0.44 |

### Beta Diversity

Dispersion (beta diversity) was similar for both the active and present taxa (Fig 3A, Table 2), and was consistent across the periods surrounding dormancy. However, the time point before corals went into quiescence (T3, 11 Dec 2020) showed significantly lower variability compared to all the other time periods, based on a Tukey HSD post hoc test (p < 0.05), and this was consistent in both the present and active microbiomes (Fig 3B, Table 2: Betadispersion).

**Figure 3.**
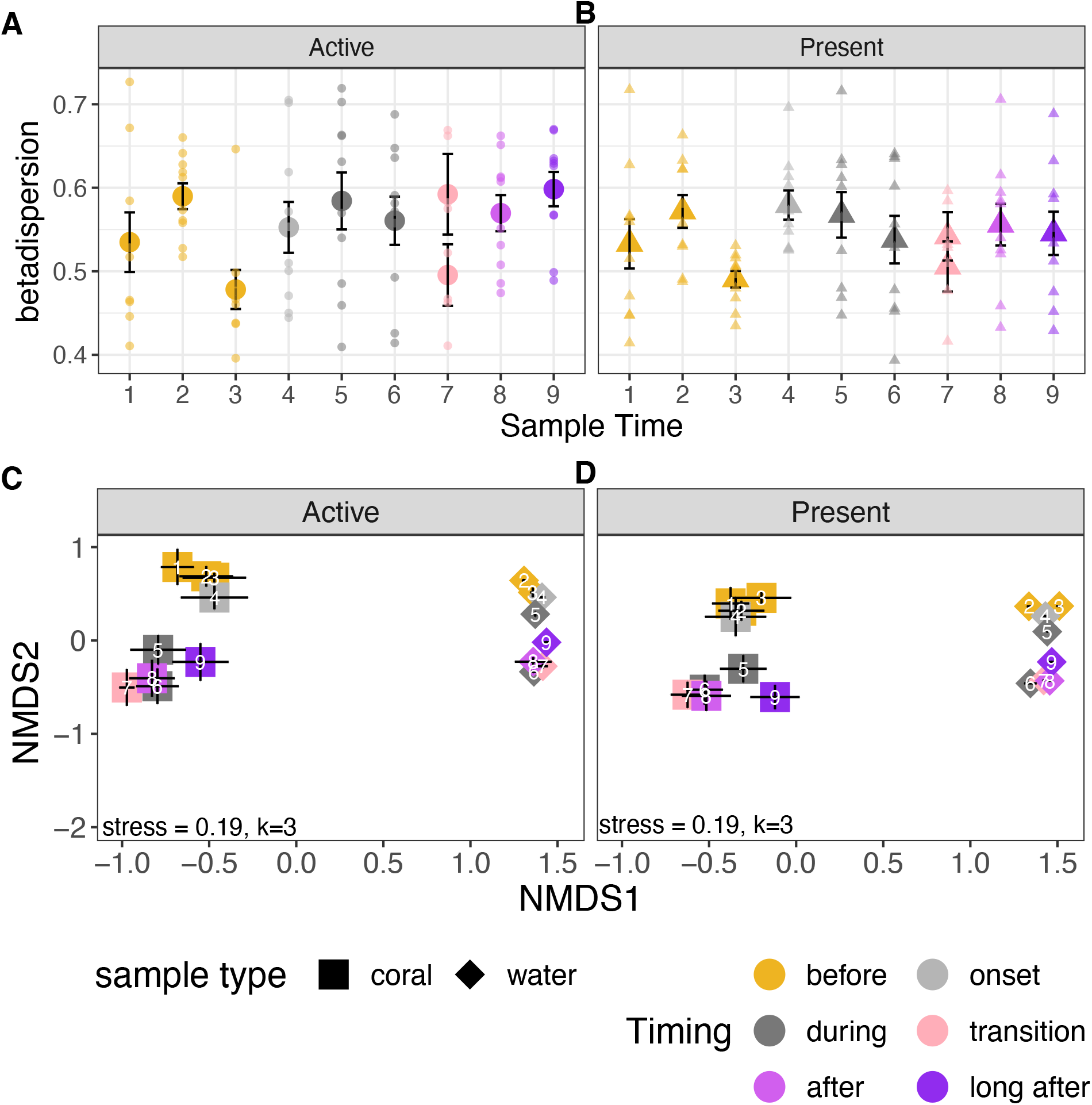
Coral microbiome beta diversity alters during quiescence. The plots show mean ± se beta dispersion of the (A) active and (B) present microbiomes for the coral (n=9-10); and mean ± se of the coral (n=9=10) and water (n=4) microbial communities in nMDS space in the (C) active and (D) present microbiomes. Colors represent timing, and shape indicates coral (square) or water (diamond) in the NMDS plot, and circle (active) and triangle (present) in the dispersion plots. Number inside of the shapes (C and D) indicate the sampling time (see Table 1). Beta dispersion (A and B) was generally high, but was significantly lowered at time point 3, the sampling point before the onset of quiescence. Microbial community composition differed significantly (p < 0.05) based on timing, sampling time and active/present microbiomes based on PERMANOVAs on the corals; and the water (plots C and D).

### Compositional shifts surrounding dormancy

*Astrangia poculata* microbial community composition shifted significantly as corals went into quiescence, and did not return to the same community after corals emerged from quiescence (Fig 3B), suggesting a reshuffling of the microbiome that persisted even two months after corals were out of quiescence. Based on the PERMANOVA analysis, we found significant effects of time (R^2^ = 0.05, p < 0.001), dormancy (before, during, after: R^2^= 0.08, p < 0.001), sample type (water/coral: R^2^ = 0.15, p<0.001), and active/present microbiomes (DNA/cDNA: R^2^= 0.02. p<0.001).

**Figure 4.**
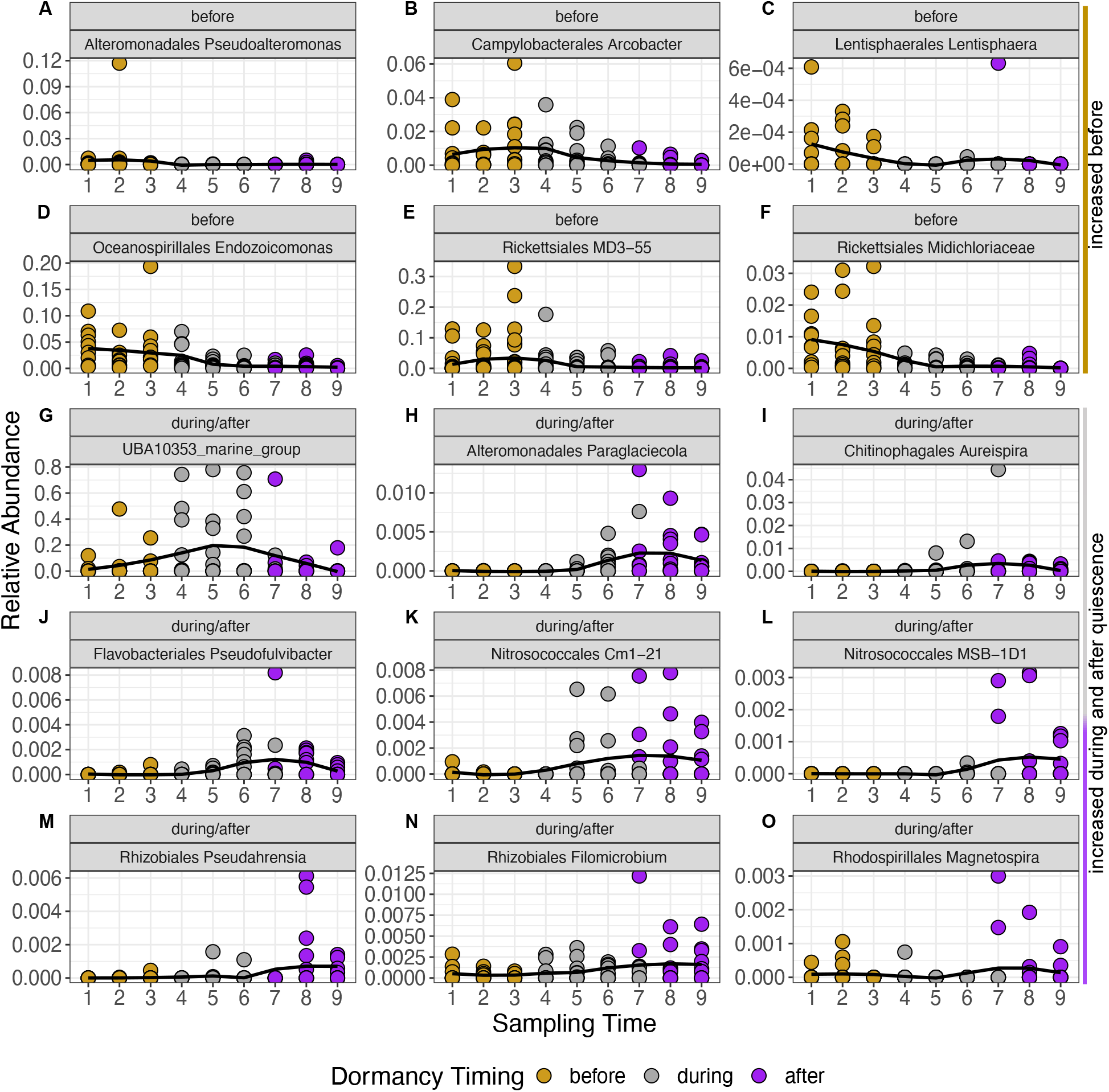
Relative abundance of selected active taxa (indicated by order and genus) that were significantly different according to the corncob analysis on the active microbiome-based ASVs. Points represent relative abundance of ASVs in each coral (9-10 samples). Lines are a loess representation of the shape of the relationship based on the geom_smooth function in ggplot (in R). Plots include taxa that are enriched before quiescence and at the first one or two time points during quiescence (A) *Pseudoalteromonas* (Altermonadales), (B) *Arcobacter* (Campylobacterales), (C) *Lentisphaera* (Lentisphaerales), (D) *Endozoicomonas* (Oceanospirillales), (E) MD3-55 (Rickettsiales), (F) Midichloriaceae. Those that were enriched during quiescence (G) UBA10353 marine group. Lastly, those that were enriched as corals were midway through quiescence and continued to increase after quiescence (H) *Paraglaciecola* (Alteromonadales),(I) *Aureispira* (Chitinophagales), (J) *Pseudofulvibacter* (Flavobacteriales), (K) Cm1-21 (Nitrosococcales). Or those that increase as corals come out quiescence (L) MSB-1D1 (Nitrosococcales), (M) *Pseudahrensia* (Rhizobiales), (N) *Filomicrobium* (Rhizobiales), (O) *Magnetospira* (Rhodospirillales). Additional data are presented in SFig1.

Both the active (cDNA) and present (DNA) microbiomes changed in similar ways in NMDS space (Figure 3B), and we did not observe significant differences in microbial community composition within a time point between the active and present communities (Table S1: Pairwise ADONIS results).

In both active and present microbiomes, communities shifted markedly between before quiescence and during/after quiescence (time points 1-3 were different from time points 5-9) (Fig 3B). The microbial community associated with the first time point for corals in quiescence (4) was not significantly different from the time points before quiescence (1-3), and from the time point one month later (5), but it differed from all future time points (6-9) (Fig 3B; Table S1). Interestingly, post-quiescent active microbiomes (time points 7, 8 and 9) did not significantly differ from corals in quiescence (time points 5 and 6) (Fig 3B). Present microbiomes showed the same pattern, except time point 9 (two months after quiescence) was significantly different from the rest of the time points (Fig 3B).

Seawater microbial community composition also changed over time significantly (R^2^ = 0.72, p =0.001) and there were significant differences in active/present seawater microbiomes (R^2^= 0.12, p = 0.001. The seawater microbiome was also consistently different from the coral microbiomes (Figure 3B).

### Dormancy associated taxa shifts

Sixty-one ASVs were identified to change significantly during the phases of quiescence in the active microbiome (Figure S1). Several taxa that were higher in abundance before corals went into quiescence began to wane at the beginning of quiescence (time points 4 and 5). These taxa include *Endozoicomonas, Arcobacter,* two groups of Rickettsiales and *Pseudoalteromonas* (Figure 4A-F). During quiescence, UBA10353 marine group showed a marked increase that lasted throughout the quiescent period (time points 4-6, Fig 4G). In late quiescence and as corals began to emerge (time points 5-9), several taxa are enriched including in orders Nitrosoccales (Figure 4K, L: Cm1-21, MSB-1D1) and Rhizobiales (Figure 4M, N), and genus *Magnetospira* (Fig 4O, for a full list, Fig S1).

Enrichment in the total present microbiome followed a similar pattern at the order level, however, it included more ASVs that shifted in abundance and presence (a total of 126, Fig S1), particularly in the orders Flavobacteriales, Chitinophagales, Cellvibrionales, and Sphingomonadales after corals emerged from quiescence and during quiescence, compared to before corals went into quiescence. Additionally, an ASV of *Candidatus* Nitrosopumilus, a taxon frequently associated with *A. poculata* (Sharp *et al*., 2017; Bent *et al*., 2021), was enriched during and after corals came out of quiescence.

Some of these taxonomic changes are likely temperature or environmentally driven, as the same taxa in the water column similarly shift in abundance (e.g., *Synechococcus,* Figure S1). However, most of the significant taxa shifts in the coral were not observed in the water column (Figure S1).

### Core microbiome

Across all time points, there were no taxa that were consistently present in the active microbiome of corals (100% of all samples), and only 5 ASVs were consistently present in 80% of samples across all time points *(Bacteroidea,* UBA4486, Terasakiellaceae, *Endozoicomonas).* Otherwise, core taxa changed by time point and varied between 4-8 ASVs within a time point, all of which were identified as significantly changing across dormancy time period with the corncob results (Figure S2).

In the present microbiome, nine taxa were present in 80% of the samples (across all time periods). These taxa include those also found in the active microbiome and *Persicirhabdus,* Pirellulaceae and *Rubripirellula.* Within a time point, core ASVs varied from 6-17 and included many that that changed significantly in the corncob results (Figure S2).

We consistently observed two archaeal ASVs in the present core microbiome, and in 65% of all active microbiomes which were associated with *“Candidatus* Nitrosopumilus” (Figure S3). Only one of these ASVs was present in the water, and only in pre-dormancy time points (Figure S3). The other *Nitrosopumilus* ASVs we observed on the corals were in low relative abundances or not detectable in any seawater samples.

### Potential functional changes

Of the 156 ASVs associated with significant shifts in the present and active microbiomes based on the corncob results, 106 taxa were assigned functions from the FAPROTAX database. Based on the taxa that were identified as significantly enriched by the corncob results and their assignment of function with FAPROTAX, we found a reduction in the number of ASVs associated with photoautotrophy and photoheterotrophy on corals (in the cDNA and DNA) as coral went into and emerged from dormancy (Figure S4A and B). We also observed a decline in the ASVs associated with intracellular parasites, nitrate reduction, sulfur/sulfite respiration and methylotrophy in the coral active and present microbiomes as corals went into dormancy. ASVs associated with nitrogen fixation, nitrification and dark sulfur oxidation increased after quiescence in the coral active microbiome.

## Discussion

Here, we show evidence from a high-resolution time series that a 3-month period of quiescence and cessation of feeding, is associated with a decrease in microbial diversity and a re-shuffling of the *Astrangia poculata* microbial community. This alteration in the microbial community persists after corals emerge from their dormant state. In particular, taxa belonging to the *Ca.* Nitrosopumilus and Nitrosococcales, groups of ammonia-oxidizing archaea and bacteria, respectively, and predicted nitrification pathways were enriched during and after quiescence. Copiotrophic bacteria as well as Rickettsiales, a proposed pathogen of corals, decreased during quiescence. This study suggests that key microbial groups and potential functions are related to quiescence in *A. poculata,* which may play an important role in this yearly dormancy period and contribute to overall holobiont health and physiology.

### Streamlining of the active microbiome diversity during dormancy, but maintenance of variability

Quiescence was associated with a streamlining of the coral’s microbiome. This was particularly evident in the active microbiome, as alpha diversity declined when corals entered dormancy. The loss of diversity is likely associated with the shedding of taxa associated with the dormancy period. Lowered alpha diversity is a consistent characteristic of host-microbiome interactions, as diapausing parasitoid wasps (Dittmer and Brucker, 2021) and ground squirrels also show a reduction in diversity of their microbiomes in dormant states (Carey and Assadi-Porter, 2017).

As corals emerged from dormancy, we observed an increase in diversity (Hill numbers D^0^, D^1^, D^2^) in both the active and present microbiome. We expect one of the drivers of this increase in alpha diversity post quiescence is the increase in feeding by the corals (Grace, 2017), thus greater exposure to externally provisioned microbes. Emergence from quiescence could also be associated with increases in colonization of microbes from the water column, leading to the observed increases in alpha diversity. Interestingly, in time point 7, when 50% of the colonies had emerged from quiescence, the corals that had already emerged from quiescence had higher alpha diversity than the corals still in quiescence, suggesting that quiescence directly influences the observed increased diversity. Further experimental work is needed to understand how quiescence emergence (e.g., mucus production, colonization from the water column) versus initiation of feeding influences the increase in diversity on corals emerging from quiescence.

Unexpectedly, coral microbiomes exhibited consistent levels of dispersion throughout the quiescence period, in both the active and present microbial communities. This was surprising, as tropical corals, and other animal hosts exposed to stressors, often show increased variability during a stressor event (the Anna Karenina Hypothesis, (Zaneveld *et al*., 2017, McDevitt-Irwin *et al*., 2017). The lowered dispersion *before* quiescence is in contrast to data that had been previously collected seasonally, in which spring collected (e.g., after quiescence) corals show decreased variability in their microbiomes relative to other seasonal timepoints (Sharp *et al*., 2017). We suggest this difference could be due to variances in the locations sampled (Rhode Island vs Massachusetts), the lower resolution of sampling timing in the previous study, or idiosyncratic differences in the environment associated with the day of sampling for the Rhode Island corals. Here, our sampling times encompassed multiple time points including those in which some corals were in quiescence and some were not (T7), one week after corals had emerged (T8) and two months later (T9), revealing that post-quiescence is not always associated with lowered dispersion, and consistently levels of betadiversity (inter-colony variability) may be an evolved trait of these corals.

### Reshuffling of the microbiome during dormancy

Quiescence was associated with a reshuffling of the coral microbial community that persisted after the corals emerged from quiescence. In fact, few taxa were consistently associated with these corals, because of the marked compositional shift that began during quiescence. The taxonomic shifts we observed before and after quiescence are similar to those observed in fall and spring collected corals (Sharp *et al*., 2017). This concurrence suggests there may be predictable or cyclical patterns in the microbiome composition associated with dormancy timing that could be important for coral holobiont health.

Among the hypotheses about the roles that microbiome plays in dormancy, the start of dormancy and how dormancy modulates the microbiome are: 1) compositional shifts that lead to the removal of pathogens, and/or 2) the replacement or maintenance of critical functions (Mushegian and Tougeron, 2019). Here, we see evidence for these two hypotheses based on the identity of the taxa and predicted metagenomes.

### Shedding of copiotrophs, including putative pathogens, during dormancy

Among the taxa that are associated with community composition shifts are the loss of copiotrophic, and particularly pathogen-associated, bacteria as corals undergo quiescence. These taxa include *Arcobacter, Pseudomonas,* and taxa in the order Rickettsiales, including a taxon that’s 97.6% identical to tropical coral parasite, *Ca.* Aquarickettsia rohweri (Klinges *et al*., 2019). Indeed, many of these taxa are associated with diseases in tropical corals (Meyer *et al*., 2019; Klinges *et al*., 2020; Becker *et al*., 2021). We also generally observed a decrease in copiotrophs, including *Endozoicomonas,* a putative beneficial symbiont in many hosts (Neave *et al*., 2016). In tropical corals, *Endozoicomonas* decreases in response to thermal stress (Lee *et al*., 2015), suggesting that it may be released when a host is stressed (e.g., an adaptive response, Buddemeier and Fautin, 1993). Because *Endozoicomonas* tends to have large genomes (4-6Mb, (Tandon *et al*., 2020), they are likely energetically costly to maintain in symbiosis (Lane and Martin, 2010), which may be why they are reduced during dormancy (and other stressor events).

We suggest the loss of copiotrophic bacteria and putative pathogens associated with the beginning of dormancy is potentially a mechanism or a consequence of a period with limited resources, when the holobiont cannot support energetically costly microbes. Thus, this loss is either the result of these microbes voluntarily or passively leaving the coral’s microbiome, or an active ejection by the coral. Alternatively, a decline in *A. poculata* holobiont metabolism may trigger a concomitant decrease in production of molecules that enrich for specific bacterial associates. Interestingly, the lone dormancy-only associated microbe was most closely related to UBA10353, a bacterial group that produces pederin, a bioactive polyketide, in sponges (Rust *et al*., 2020). A resulting hypothesis is that the increase in this bacterium could result in production of antimicrobial compounds to reduce colonization of microbes while the host is quiescent, and/or help explain the loss of microbes associated with dormancy. In contrast with previous conclusions from seasonal characterization of *A. poculata* microbiomes (Sharp *et al*., 2017), the higher resolution sampling in this study reveals that putative pathogens are not higher in proportional abundances in winter months, as previously described (2017), but rather are at their maximum just before entry into quiescence, and are then shed after initiation of quiescence.

### Microbes involved in essential functioning during dormancy

During dormancy we observed an increase and maintenance of microbes associated with ammonia oxidation, nitrification, and nitrogen fixation, suggesting the microbiome plays a role in the maintenance and acquisition of nitrogen while corals are not actively feeding. Among the taxa that likely contribute to replacing host functions, were archaea in the genus, *Ca.* Nitrosopumilus, known associate of *Astrangia poculata* (Sharp *et al*., 2017; Bent *et al*., 2021); and the bacteria in the order Nitrosococcales, (Cm1-21, MSB-1D1).

*Ca.* Nitrosopumilus and Nitrosococcales are common ammonia oxidizers ((Semedo *et al*., 2021). Here, corals are not actively feeding and do not have any visible algal symbionts, thus these ammonia oxidizers may play an important role in nitrate acquisition for the host, or other essential microbes. Corals in late quiescence and early non-quiescence states also show increases in nitrate-reducing Rhizobiales *(Psuedahrensia* and *Filomicrobium,* (Jung *et al*., 2012), and *Magnetospira,* a likely nitrogen fixer (Williams *et al*., 2012). These taxa, along with the ammonia oxidizers suggest the microbial community likely continues to bolster nitrogen cycling in the host as corals emerge from quiescence, and may help build energetic reserves that were depleted during quiescence (Trumbauer *et al*., 2021). The presence of these bacteria and archaea, particularly the ammonia oxidizers, during dormancy and after dormancy may help explain some of the acquisition of nitrogen (i.e., ammonia, the host’s preferred DIN source) for the host in general, which is usually attributed to heterotrophy and enhanced by algal symbionts (DiRoberts *et al*., 2021; Trumbauer *et al*., 2021). For example, the increase in nitrifying microbes may explain the higher δ^15^N values in *A. poculata* tissues previously found in the winter compared to the fall, (Trumbauer, Grace, and Rodrigues 2021) as winter corals are quiescent and are not relying on heterotrophy (or photoautrophy). More research is needed to understand the role of the microbiome in the cycling of nitrogen in coral tissues, and how this may impact coral fitness after quiescence.

Corals in late quiescence and after emergence also showed increases in Flavobacteriales (e.g., Pseudofulvibacter, Ulvibacter, Figure 4, Figure S1, S4). As microbial heterotrophs, it is possible that these taxa are playing a role in carbon cycling on and/or with the host before and just as the coral begins to feed actively. In sponges, for example, those with high heterotrophic microbial loads, show that microbes play a role in DOM assimilation (Rix *et al*., 2020). Alternatively, the increase in Flavobacteriales may be due to increases in food availability and are not necessarily host-associated (Gavriilidou *et al*., 2020).

Replacement of host nutrition during dormancy is a common theme in host-microbial systems. In ground squirrels *(Ictidomys tridecemlineatus),* the gut microbiome plays a critical role during hibernation to recycle nitrogen (from urea), which supports tissue growth while the animal is not feeding (Regan *et al*., 2022), which is evolutionary advantageous leading up to the breeding season. Furthermore, diapausing *Daphnia* eggs are enriched in *Nitrospira* bacteria, suggesting nitrification may be a function of dormant microbiomes in other hosts (Mushegian *et al*., 2018). Nutritional provisioning, particularly of nitrogen, during dormancy is potentially a convergent trait across host-microbes that undergo dormant periods.

### Onset and Emergence from Quiescence Timing

Predictable, seasonal quiescence in *A. poculata* has been observed and documented in the field (Grace, 2017) and in the lab, quiescence can be experimentally triggered by lowering temperatures to 5°C (Grace, pers comm, (Wuitchik *et al*., 2021). Emergence from dormancy was similarly found by raising temperatures above 5°C (Wuitchik et al. 2021). Here, our findings support that the onset of dormancy was associated with temperatures reaching 5°C, suggesting winter temperature as one of the environmental triggers for the onset of dormancy. Indeed, corals experienced temperatures as low as 2°C (Figure 1). However, absolute temperature is likely not the only trigger for emergence: corals began to emerge from dormancy while water temperatures remained close to 5°C (Fig 1, Table 1), thus there are likely other factors that lead to the cessation of quiescence. As the microbiome shifts swiftly during quiescence, it is possible there is an interplay between the environment, the host, and the microbiome that triggers the onset and emergence from quiescence. Further experimental work, including isolating the effects of temperature and season, and characterizing the mucus metabolome throughout seasonal and environmental shifts, will help to elucidate the relationship between microbial shifts and quiescence. Because *A. poculata* are facultatively symbiotic with microalgae, and can also be used in aquaria-based studies (Bent *et al*., 2021) it is an ideal marine experimental system for investigation of the role of the microbiome in animal host dormancy.

### Conclusions

Our findings suggest that *A. poculata* quiescence is involved in re-shuffling the microbiome, leading to a new, persistent microbiome community structure. This shuffling is associated with shedding potential pathogens, like Rickettsiales, and a shift in the taxa that likely replace nutrition, like *Ca.* Nitrosopumulis, while the host is inactive (Figure 5). Overall, this study demonstrates that key microbial groups are related to quiescence in *A. poculata* and may play an indirect or direct role in onset and emergence from dormancy. Further understanding of the interactions between the coral and these specific microorganisms during this change in coral metabolic status will advance our understanding of coral host-microbiome dynamics and dormancy.

**Figure 5.**
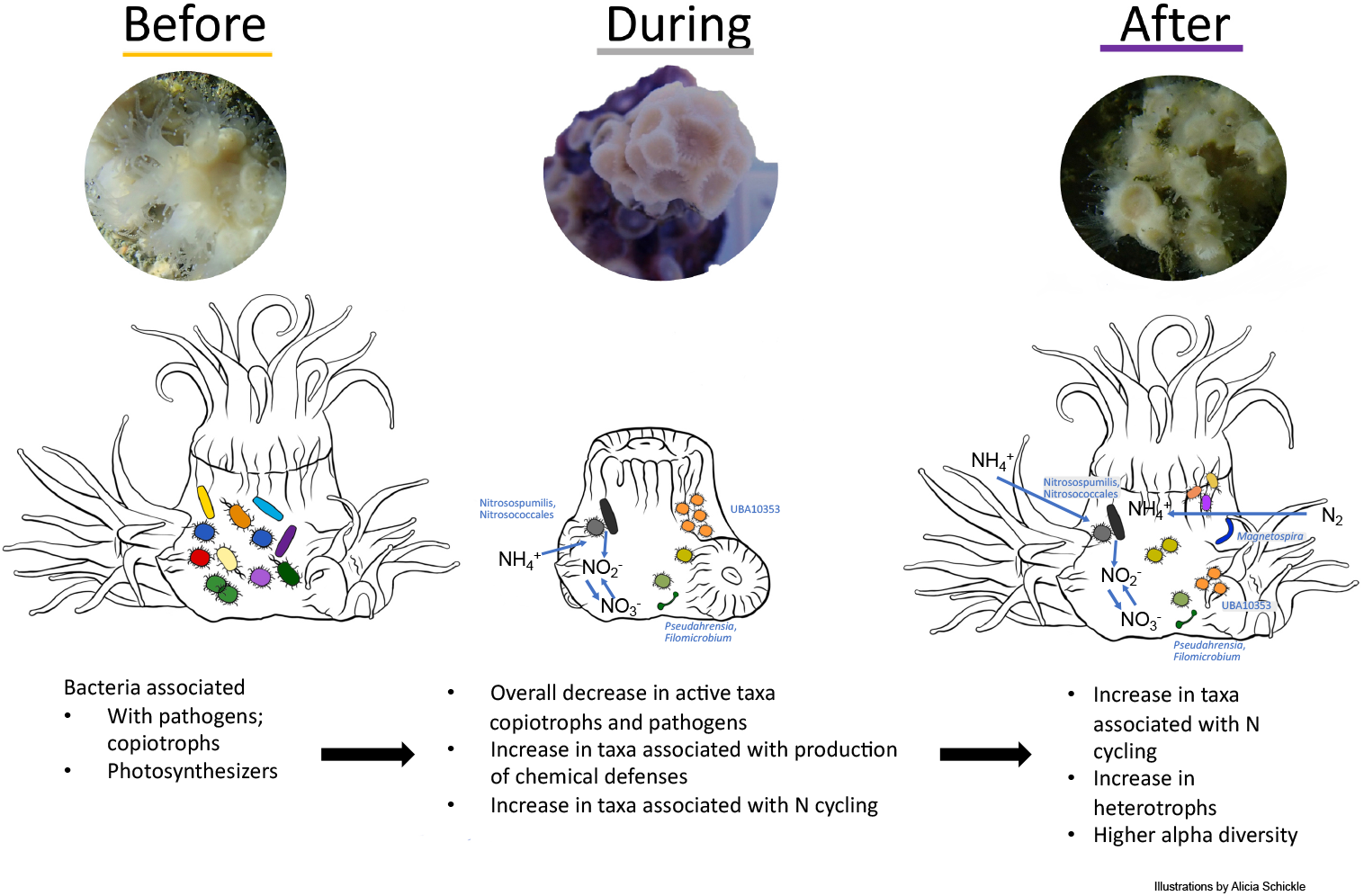
Conceptual diagram of the diversity and community shifts that occurred in the *Astrangia poculata* microbiome before, during and after quiescence. Illustrations by: Alicia Schickle

## Acknowledgements

We would like to thank Sean Grace for his advice and input on quiescence, what to look for and when to sample. We would also like to thank the WHOI dive team, Kim Malkowski and Ed O’Brien for dive assistance and coral monitoring. Thanks to the Apprill lab, particularly C. Becker for help with a critical time point. A.L. Brown was funded by a WHOI Postdoctoral Scholar Award. Additional funding support included a NOAA OAR Cooperature Institutes (#NA19OAR4320074) and National Science Foundation award (OCE-1938147) to AA. Additional support included funded awarded to KS, via the Institutional Development Award (IDeA) Network for Biomedical Research Excellence from the National Institute of General Medical Sciences of the National Institutes of Health under grant number P20GM103430. The authors extend appreciation to the annual Temperate Coral Research Conferences hosted by Roger Williams University, Boston University, and Southern Connecticut State University, for fostering creative conversations and collaborations leading to this work.

## Supplemental Figures

Figure S1) Differential abundance of ASVs (each point) Before vs During and Before vs After quiescence in the coral (active and present) and water (active and present). Values are effect size ± se output from the corncob model for each ASV. Points are colored based on whether they were enriched before (coral: yellow, water: light blue), during (coral: gray, water: blue), and after (coral: purple, water: dark blue).

Figure S2) Relative abundance of ASVs associated with the core microbiome (at 80% prevalence) of the cDNA and DNA. Each facet is labeled by Order and Genus. Points are colored by sample type (yellow indicates coral, blue indicates water) and present/active microbiomes (lighter colors represent the active microbiome and darker colors represent the present). Lines are created by the loess function in ggplot2.

Figure S3. Relative abundance across all replicates in a time points of ASVs in the archaeon genus *Ca.* Nitrosopumilus in the coral (A) active, (b) present microbiomes and the water (c) active and (d) present microbiomes for each sampling point. Each color within the bar represents a different ASV associated with the DNA core (at 80% prevalence) or cDNA core (at prevalence 65%), represented by the asterisk (ASV 2,3) or ASVs that were significantly different based on the corncob results represented by ^‡^ (ASV 1 and 2). ASV 2 was significantly differentially abundant in the DNA based on the corncob results and is part of the coral core community (Fig S2).

Figure S4. Potential functions based on the FAPROTAX database (Liang *et al*., 2020) assigned to the ASVs (here represented at the genus level) that are significantly changing based on the corncob results. A point represents that the function is present in that genus in the (A) present microbiome (i) before versus after corals are in quiescence and (ii) before versus during corals are in quiescence; and (B) in the active microbiome (i) before versus after corals are in quiescence and (ii) before versus during corals are in quiescence. Colors represent if the taxa/function are enriched before, during or after quiescence.

